# Momelotinib is a highly potent inhibitor of FLT3-mutant AML

**DOI:** 10.1101/2021.03.18.436047

**Authors:** Mohammad Azhar, Zachary Kincaid, Meenu Kesarwani, Tahir Latif, Daniel Starczynowski, Mohammad Azam

**Affiliations:** Division of Pathology, Cincinnati Children’s Hospital, Cincinnati, Ohio; Department of Pediatrics, University of Cincinnati, Cincinnati, OH; Division of Experimental Hematology and Cancer Biology, Cincinnati Children’s Hospital, Cincinnati, OH; Department of Internal Medicine University of Cincinnati and Director of Infusion Services; Department of Cancer Biology, University of Cincinnati, Cincinnati OH

## Abstract

Kinase activating mutation in FLT3 is the most frequent genetic lesion associated with poor prognosis in acute myeloid leukemia (AML). Therapeutic response to FLT3 tyrosine kinase inhibitor (TKI) therapy is dismal, and many patients relapse even after allogenic stem cell transplantation. Despite the introduction of more selective FLT3 inhibitors, remissions are short-lived, and patients show progressive disease after an initial response. Acquisition of resistance-conferring genetic mutations and growth factor signaling are two principal mechanisms that drive relapse. FLT3 inhibitors targeting both escape mechanisms could lead to a more profound and lasting clinical responses. Here we show that the JAK2 inhibitor, momelotinib, is an equipotent type-1 FLT3 inhibitor. Momelotinib showed potent inhibitory activity on both mouse and human cells expressing FLT3-ITD, including clinically relevant resistant mutations within the activation loop at residues, D835, D839, and Y842. Additionally, momelotinib efficiently suppressed the resistance mediated by FLT3 ligand (FL) and hematopoietic cytokine activated JAK2 signaling. Interestingly, unlike gilteritinib, momelotinib inhibits the expression of MYC in leukemic cells. Consequently, concomitant inhibition of FLT3 and downregulation of MYC by momelotinib treatment showed better efficacy in suppressing the leukemia in a preclinical murine model of AML. Altogether, these data provide evidence that momelotinib is an effective type-1 dual JAK2/FLT3 inhibitor and may offer an alternative to gilteritinib. Its ability to impede the resistance conferred by growth factor signaling and activation loop mutants suggests that momelotinib treatment could provide a deeper and durable response; thus, warrants its clinical evaluation.

**Key points:**

1. Momelotinib shows high efficacy against FLT3 mutated AML cells including quizartinib-resistant activation loop variants.
2. Momelotinib effectively suppresses intrinsic resistance conferred by FLT3 ligand and hematopoietic cytokines (GM-CSF and IL3).

## Introduction

Kinase activating mutations in the *FMS-like tyrosine kinase 3* (*FLT3*) gene represent the most frequent molecular lesion in acute myeloid leukemia (AML)^1-3^. Approximately one-third of AML patients harbor internal tandem duplication (ITD) of the juxta-membrane region of FLT3, which is associated with poor treatment outcome and overall survival even after stem cell transplantation^4,5^. Additionally, a significant number of patients (7%) reported to have kinase activating mutations from the activation loop (D835) with an unknown prognosis^6,7^. Several small-molecule FLT3 tyrosine kinase inhibitors (TKIs) have been evaluated in the last two decades, but none could induce a durable response. Because monotherapy of first-generation FLT3 inhibitors (midostaurin, lestaurtinib, and sorafenib) failed to exert expected clinical response, a combination with conventional chemotherapies were sought^8^. Only midostaurin demonstrated modest clinical benefit in the combination that resulted in its approval for clinical use in 2017^9,10^.

Second generation FLT3 inhibitors, quizartinib, gilteritinib, and crenolanib, were designed for greater selectivity and specificity^11^. They showed a narrow kinome-profile and better clinical efficacy than first-generation inhibitors but failed to induce a durable response. For instance, like imatinib, quizartinib, a type-II inhibitor (binds to inactive conformation), is prone to select kinase activating resistant mutations from the gatekeeper and activation loop residues^12,13^. Therefore, patients harboring FLT3-TKD mutations (D835) are non-responsive to quizartinib. Besides, its activity against the c-Kit receptor causes significant myelosuppression. In contrast, gilteritinib and crenolanib are type-I inhibitors (bind to active conformation) that can inhibit both FLT3-ITD and FLT3-TKD and efficiently suppress the secondary drug-resistant mutants except the gatekeeper variant, F691L^14^. Unlike quizartinib, both gilteritinib and crenolanib do not inhibit c-KIT. Therefore, they are not myelosuppressive, and patients show better hematologic recovery ^3,15,16^. However, the median duration of response remains short-lived, weeks to months, and most patients develop resistance by acquiring mutations in other signaling pathways, such as RAS, PTPN11, TET2, and IDH1/2^17,18^. Moreover, patients treated with gilteritinib display QT prolongation, likely due to on-target inhibition of hERG^19^. FLT3-ITD positive leukemic cells in BM are refractory to TKI inhibitors, which serve as a reservoir to develop resistance. FLT3 ligand (FL) ^20^, chemokines, and cytokines (CXCR4, FGF2^21^, IL-3 and GM-CSF^22^) from the stroma confer TKI refractoriness. While cytokine (GMCSF and IL-3) mediated resistance mediated by JAK2 activation, which can be tackled by combined FLT3 and JAK2 inhibition^22^. However, the mechanisms of FL driven TKI resistance is not clear. Nonetheless, activation of FLT3-WT signaling by FL was implicated in averting the TKI response^23^. Altogether, growth factor signaling abrogates TKI response, and failure to completely eradicate leukemic cells results in minimal residual disease (MRD).

AML patients who achieved complete remission lacking FLT3-ITD clones showed better overall survival than patients with measurable MRD, suggesting that eradicating the FLT3-ITD clones will have a deeper response with better overall survival. Thus, an ideal FLT3 inhibitor should be able to suppress FLT3-ITD, FLT3-TKD, JAK2, FL-mediated resistance but and sparing c-KIT receptor and hERG to avoid myelosuppression and cardiotoxicity due to QT prolongation. Here we show that Momelotinib, a JAK2 inhibitor^24^, is on fast-track designation by the FDA for myelofibrosis treatment, is an equi-potent type-I FLT3 inhibitor. Like gilteritinib, it efficiently suppresses FLT3-ITD and drug-resistant FLT3 mutants from the activation loop and lacks activity against the c-KIT receptor. Thus, providing an explanation why momelotinib treatment is not myelosuppressive and alleviates anemic symptoms in myelofibrosis (MF) patients, which is commonly observed with currently used JAK2 and FLT3 inhibitors (ruxolitinib, fedratinib, and quizartinib)^25,26^. Unlike gilteritinib, momelotinib treatment did not show QT prolongation and cardiotoxicity suggesting it does not inhibit hERG^27^. Structural modeling revealed that it binds to an active conformation of FLT3 kinase, thus providing an explanation for its activity against the mutants from the activation loop. Perhaps more interestingly, in contrast to gilteritinib, it can efficiently suppress the FL and cytokine-activated Jak2 signaling mediated resistance. Moreover, unlike gilteritinib, momelotinib treatment suppresses the expression of MYC by inhibiting the IKBKE^28^. Depletion of Myc by BRD4 inhibition is under clinical evaluation in AML driven by RAS and MLL-fusion^29,30^. Given that the momelotinib has been approved for myelofibrosis treatment, our preclinical data suggest that it will be more effective in suppressing AML than the currently used FLT3 TKIs and may induce stable and prolonged remission.

## Methods

### Plasmids and inhibitors

pMSCV-Ires-GFP, pMSCV-FLT3-WT-Ires-GFP, pMSCV-FLT3-ITD-Ires-GFP were described earlier. JAK2 and ABL inhibitors were purchased from Chemietech Inc. and LC Chemicals. Quizartinjb, Gilteritinib, and Momelotinib were purchased from AdooQ Biosciences.

### Cell Lines

BaF3 cells stably expressing quizartinib resistant variants were generated as described earlier^31^. Human AML cell lines K562, Molm13, and MV4-11 were described earlier^32^.

### Cell-viability and western-blotting assays

Ten thousand BaF3 cells stably expressing Jak2-V617F, FLT3-WT, FLT3ITD, and drug-resistant variants of FLT3ITD were plated in triplicate in 96-well plates in 0.1 ml RPMI medium containing 10% FCS with varying inhibitor concentrations. According to the manufacturer’s specifications, after 60 hours, cell viability was assessed using WST-1 reagent (Roche Applied Science, USA). Absorbance, A450 nm, was averaged and plotted against inhibitor concentration as a best-fit sigmoidal curve, using the Prizm (GraphPad). The concentration resulting in 50% maximal inhibition was reported as the cellular IC50. FLT3, STAT5, and ERK1/2 immunoblotting were performed as described earlier.

### Structural modeling, inhibitor docking, and representation

Targeted docking of momelotinib to the ATP-site and blind docking to whole-kinase domain structure was performed using SwissDock^33,34^ and FLT3 kinase coordinates with gilteritinib(PDB: 6JQR) and quizartinib (4XUF). The algorithm of Swissdock generates a large number of binding modules (typically 5000 to 15,000)—either by local docking (in a user-defined box) or on the entire protein surface (blind docking), which identifies unanticipated binding sites and cavities. Simultaneously, free energies for ligand binding using CHARMM are estimated on a grid^35^ and ranked. Modules with the most favorable energies are clustered^36^, and the most favorable clusters are contained in the output PDB file. For each cluster, binding modules with the lowest energy (i.e., the most likely to represent correct binding) are selected for further verification. Figures were generated using PyMol.

### In vivo efficacy of Momelotinib

Molm13 and MV4-11 cells were transduced with lentiviruses co-expressing firefly luciferase and cherry fluorescent protein. HEK293T cells were transfected with pMSCV-Luc-cherry-GW27 to generate high titer viruses. One million cherry-positive cells were injected into the tail veins of immunocompromised NSGS mice (Jackson Laboratories, USA). Two days post-transplantation, mice were intraperitoneally administered with momelotinib (100 mg/kg/daily) for two weeks. On days 7 and 14, mice were imaged. Imaging and analysis were performed as described earlier. All experiments involving mice were performed according to the National Institutes of Health Guide for the care and use of laboratory animals and were approved by the Cincinnati Children’s Hospital Institutional Animal Care and Use Committee (IACUC).

## Results

### Activity of Momelotinib against FLT3 kinase

While studying the efficacy of combinatorial targeting of JAK2 and FLT3 using ruxolitinib or momelotinib (as JAK2 inhibitors) with quizartinib (as a Flt3 inhibitor) on AML cells, we observed inhibitory activity of momelotinib on FLT3^ITD^ cells. Unlike ruxolitinib, momelotinib alone can efficiently suppress the proliferation of both BaF3-JAK2V617F (IC50: 325 nM), BaF3-FLT3-WT (IC50: 320 nM), and BaF3-FLT3-ITD (IC50: 401 nM) (**Figure 1A** and **Supplementary Fig.1A**). Similarly, like quizartinib and gilteritinib, momelotinib inhibited the proliferation of patient-derived AML cells harboring FLT3^ITD^, MV4-1, and Molm13, with IC50s of 203 and 165 nM, respectively (Fig 1B and Supplementary Figure 1 C and D). In contrast, momelotinib failed to inhibit the proliferation of the BCR-ABL mutant human AML cell line, K562, which lacks FLT3-ITD mutation (**Figure 1B** and **Supplementary Figure 1C**,**D**). To determine whether the inhibitory effect of momelotinib on cell proliferation is mediated through on-target inhibition of FLT3 signaling, FLT3 autophosphorylation, and phosphorylation of STAT5 and ERK1/2 were analyzed by immunoblotting. Like quizartinib and gilteritinb, momelotinib led to a dose-dependent decrease of pFLT3, pSTAT5, and pERK1/2 in both BaF3-FLT3-ITD and human cell lines, Molm13 and MV4-11 (**Figure 1C,D** and **Supplementary Figure 1 B**). In contrast, the JAK family inhibitor ruxolitinib, which has no activity against FLT3 failed to inhibit phospho-FLT3 and phospho-STAT5, suggesting the inhibition of FLT3 signaling is independent of JAK2 inhibition (**Supplementary Figure 1A**,**B**). These data demonstrate that momelotinib effectively inhibits mutant FLT3 and its downstream signaling pathways.

**Figure 1.**
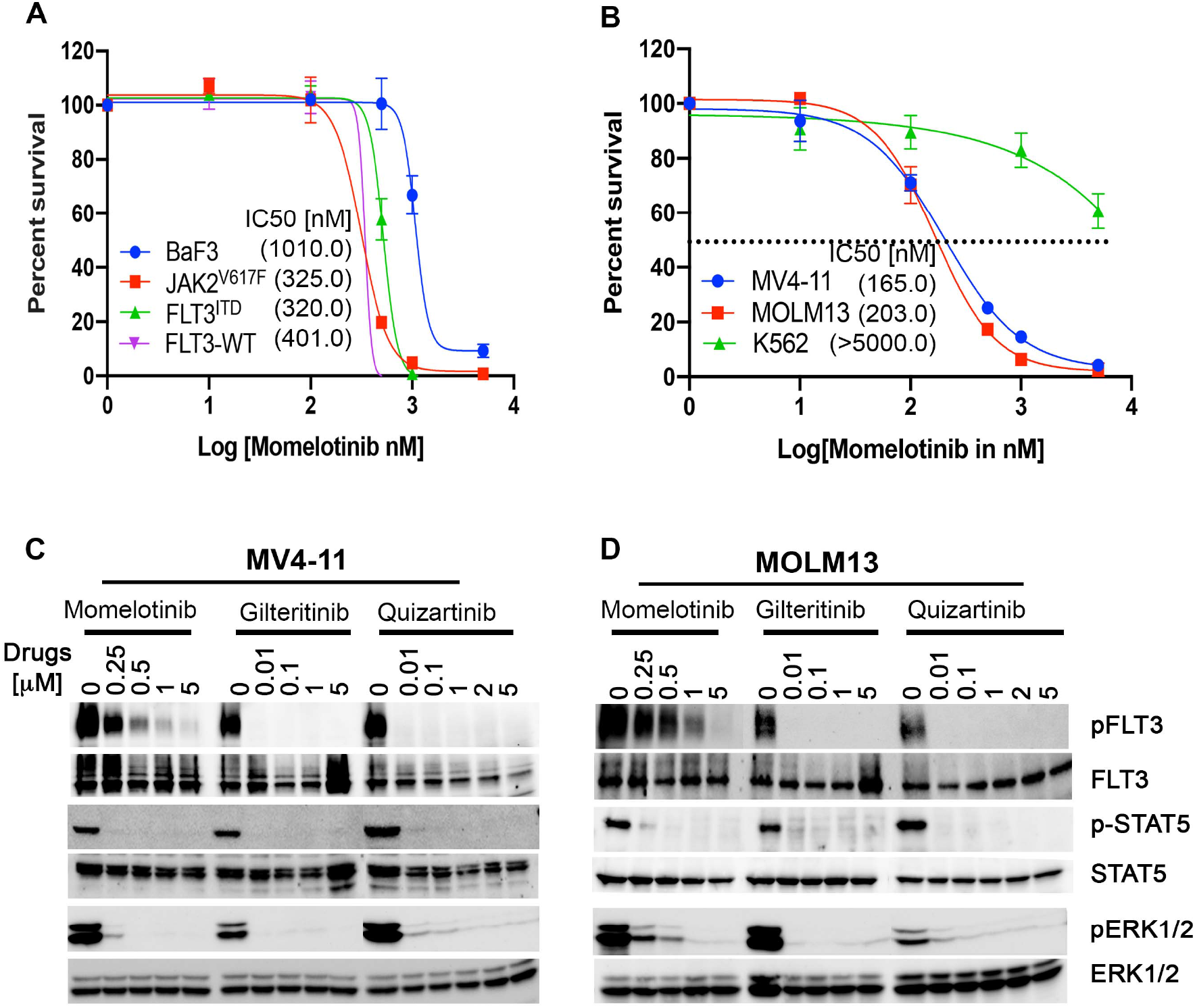
Momelotinib potently inhibits both mouse and human FLT3-ITD positive leukemic cells viability and FLT3 signaling. (**A**) Sigmoidal curve showing the viability of BaF3 cells expressing FLT3-WT, Jak2V617F and FLT3-ITD treated with DMSO or increasing concentrations of momelotinib for 72 hrs. (**B**) Sigmoidal curve showing the potent inhibition of human FLT3-ITD mutant AML cells (MV4-11 and MOLM13) while proliferation of K562 cells were not significantly affected. IC50 values for each cell line is indicated in parenthesis. (C and D) Showing pFLT3, pSTAT5 levels determined by western blotting using total cells extracts of MV4-11 (C) and MOLM13 cells (D) treated with increasing concentrations of momelotinib for two hours.

### Momelotinib suppresses the proliferation of TKI resistant variants of FLT3-ITD

Next, we sought to determine the efficacy of momelotinib against clinically relevant FLT3-ITD resistant variants. Like gilteritinb, momelotinib potently inhibited the proliferation of FLT3-ITD resistant variants D835Y, D839G, and Y842H except the gatekeeper mutant, F691L (**Fig 2 A-C**). Accordingly, western blotting of phospho-STAT5 and phospho-FLT3 showed reduced phosphorylation at concentrations of momelotinib that parallels the concentrations required to inhibit cell survival (Fig 2D). Furthermore, we tested the efficacy of momelotinib on FLT3-ITD compound mutations (combination of gatekeeper mutation with activation loop variants) to determine its clinical utility as they pose significant challenge. We generated BaF3 cells expressing FLT3-ITD-F691L with most prominent activation loop mutants (D835Y and Y842H on the same allele. Inhibitory activity of momelotinib was determine and compared with FLT3-ITD and FLT3-ITD/F691L alone. We observed increased sensitivity (∼four-fold) of compound mutant FLT3-ITD-F691L/Y842H (IC50: 1001 nM) compared to FLT3-ITD-F691L (IC50:3976 nM). In contrast, compound mutant FLT3-ITD-F691L/ D835Y did not show any significant change in IC50 value compared to FLT3-ITD-F691L. These data suggest that mutations from the activation loop may break the resistance conferred by the gatekeeper mutation F691L. It seems F691L mutation stabilize a unique conformation (SRC-like intermediate conformation) to prevent momelotinib binding rather than a steric hindrance observed with type-II inhibitor, quizartinib. Therefore, a combined F691L and Y842H mutation shifts the equilibrium to an active state conformation to which momelotinib seemingly has a greater affinity. It also suggests that compound mutation F691L/D835Y does not significantly change the conformational dynamics and, therefore unable to alter the IC50.

**Figure 2.**
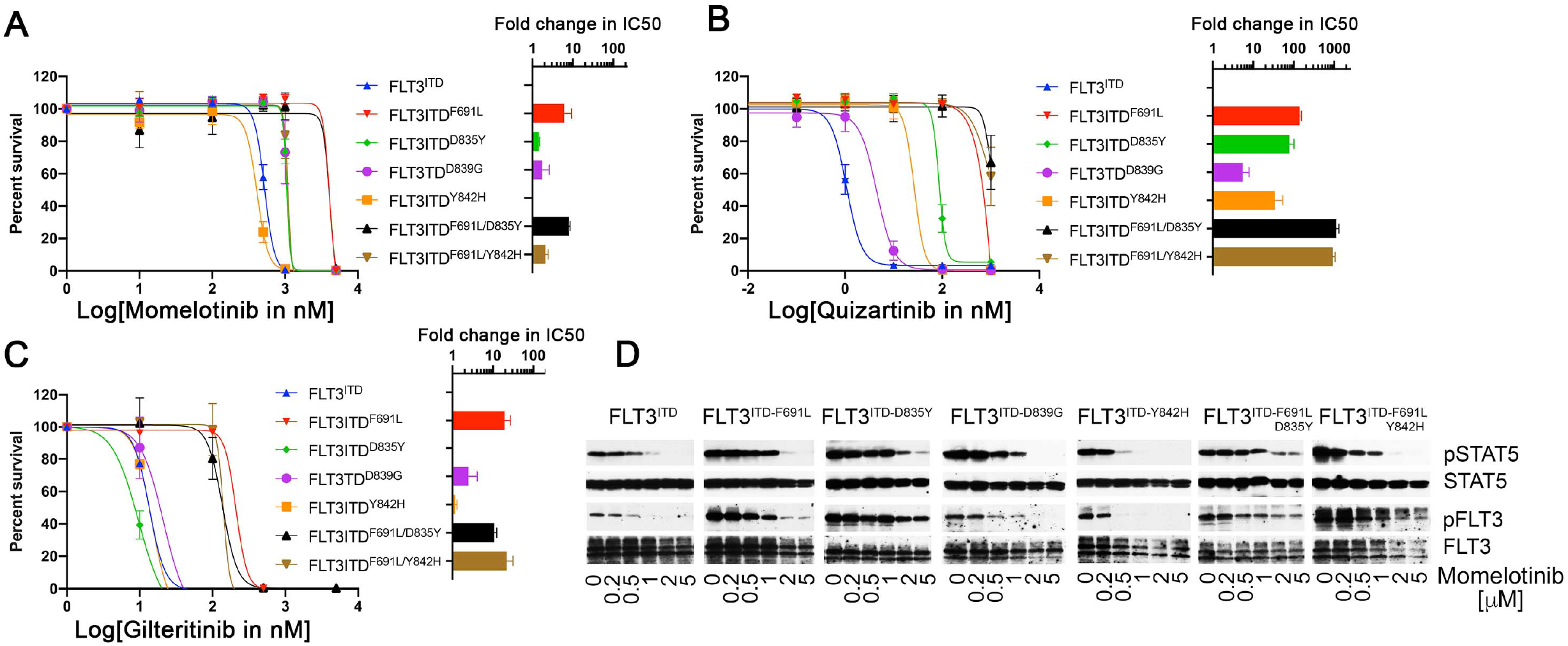
Momelotinib, like gilteritinib, efficiently inhibits the proliferation of FLT3-ITD quizartinib resistant mutants from the activation loop. (**A-C**) Dose–response sigmoidal curve showing the proliferation of BaF3 cells expressing FLT3-ITD and its kinase domain quizartinib resistant variants at different concentrations of momelotinib (A), quizartinib (**B**) and gilteritinb (**C**). Fold difference in the IC50 values for each FLT3-ITD variants normalized to FLT3-ITD are presented as a bar graph on the right in logarithmic scale. Note, momelotinib efficiently inhibits the activation loop quizartinib-resistant mutants as well as compound mutant FLT3-ITD-F691L/Y842H which is fully resistant to gilteritinib (see the brown bar). (**D**) Immunoblot analysis showing inhibition of the kinase activity of FLT3-ITD and resistant variants treated with momelotinib at different concentrations. Total cell extracts from the cells treated with momelotinib for two hours were probed with anti-phospho-FLT3, anti-phospho-STAT5, anti-FLT3 and anti-STAT5 antibodies.

### Momelotinib is a type-I FLT3 inhibitor

Next, we performed molecular docking studies of momelotinib to understand the structural basis of FLT3 inhibition. We used both inactive conformation of FLT3 with quizartinib (PDB: 4XUF) and active conformation crystallized with gilteritinib (PDB: 6JQR) and FF-10101F (PDB: 5×02). Our *in-silico* analysis failed to predict the binding of momelotinib at the ATP site when inactive conformation was used for docking. However, we observed efficient binding of momelotinib to the ATP site with the active conformation of FLT3. Our docking analysis predicts that momelotinib binds to an open and enzymatically active conformation of FLT3, in which residue Phe 830 of the DFG motif is displaced to accommodate phenyl-cyanomethyl ring (**Fig 3**). Momelotinib anchors to ATP site by three hydrogen bonds with residues Cys 694 from the hinge region and with residues Glu 817 and Ser 618 from the P-loop (**Figure 3B**). Unlike quizartinib, kinase conformation with DFG-out conformation (inactive conformation) is incompatible for both momelotinib and gilteritinb binding due to steric clash with Phe 830 (**Figure 3 D-G**) The enhanced activity of momelotinib against the A-loop variants (D835Y, D839G, and Y842H) is possibly due to the stabilization of an open and active conformation by these mutants to which it preferentially binds. Like gilteritinib, momelotinib is not in close contact with gatekeeper residue F691. A leucine substitution for phenylalanine at this residue has conferred resistance to both type-I and type-II inhibitors. However, the extent of resistance is much higher for type-II inhibitor, quizartinib, because of direct steric hindrance (**Figure 2B** and **3C, D**). It is not clear how F691L confers resistance to type-I inhibitors. Our model suggests that substitution of Leu for Phe 691 will weaken the hydrophobic spine that destabilizes the active conformation that possibly stabilizes an intermediate conformation to which type-I inhibitors seemingly have a lower affinity. This supports the notion that momelotinib activity against FLT3-ITD-F691L/Y842H is likely due to stabilization of an open and active conformation. Increased FLT3 autophosphorylation of FLT3^ITD^- F691L/D842H compared to FLT3-ITD-F691L/D835Y from the untreated extract provides evidence that stabilization of gatekeeper mutant to a fully active conformation would sensitize it to inhibition by type-I inhibitors (**Figure 2D**).

**Figure 3.**
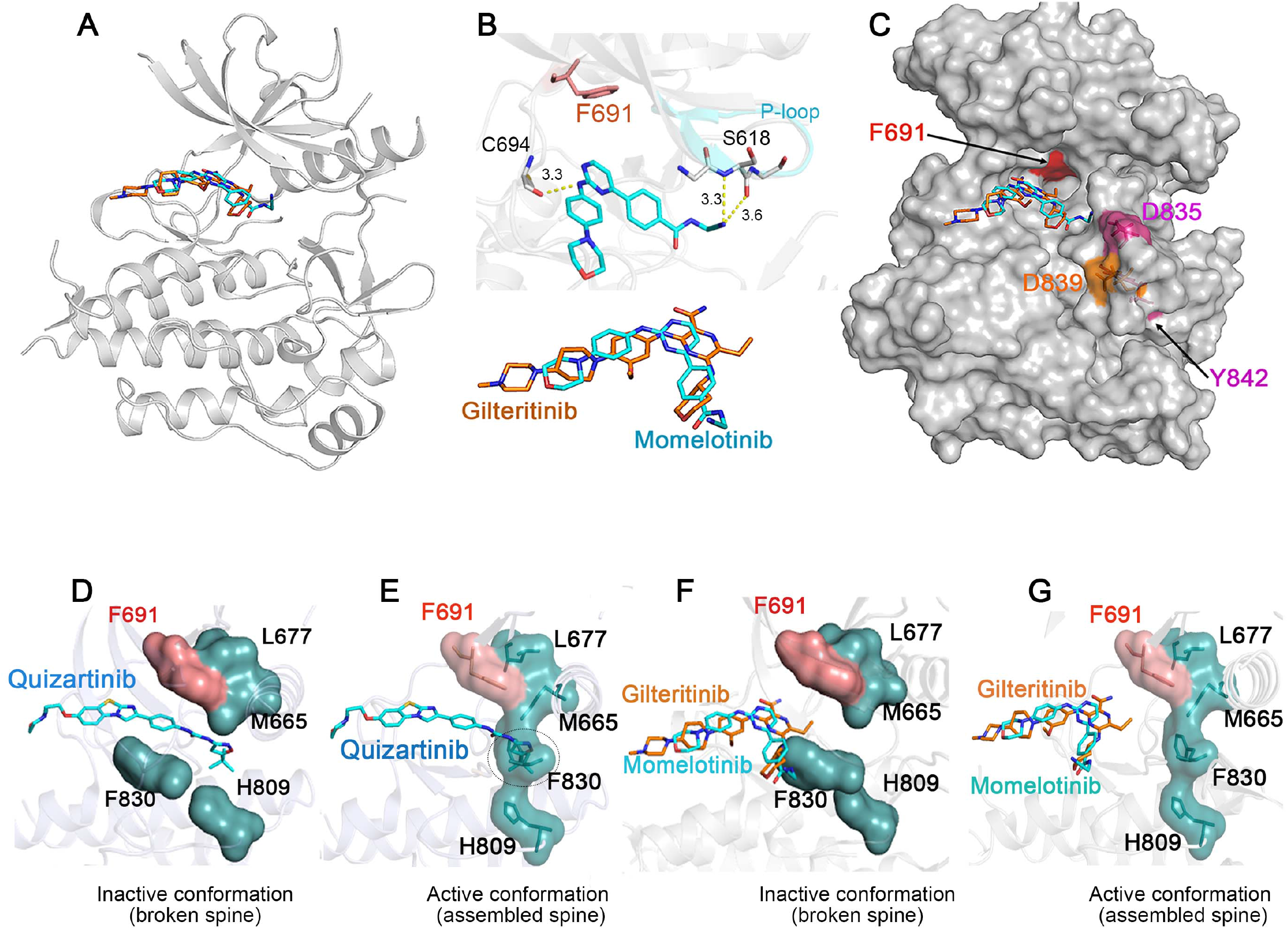
Momelotinib is a Type-I FLT3 inhibitor. (**A**) Ribbon depiction of a structural model of FLT3-momelotinib using the coordinates of the FLT3 kinase with quizartinib (PDB:4XUF) and gilteritinib (PDB:6JQR). A stick representation of momelotinib (cyan) and gilteritinib (brown) showing their binding to ATP site. (**B**) A close-up view of the active site of the FLT3 kinase showing the interaction of momelotinib with Cys 694 from the kinase hinge region and Ser 618 from the P-loop through three hydrogen bonds. Bottom panel showing a similar binding of momelotinib and gilteritinb. (**C**). A surface depiction of FLT3 kinase-with gilteritinib showing the binding of both momelotinib and gilteritinib. Activation loop residues D835 (pink), D839 (brown) and Y842 (pink) frequently mutated in patients treated with type-II inhibitors are shown. (**D**) The central para-substituted phenyl of quizartinib is sandwiched between the gatekeeper residue, Phe 691, and the phenylalanine residue in the DFG motif, Phe 830 through a π-π interactions to stabilize an inactive, type-II, conformation. (**E**) A model of active FLT3 conformation with quizartinib defined by stable hydrophobic spine showing steric clash between quizartinib and Phe 830 of DFG motif. (**F**) A model of FLT3 inactive conformation docked with momelotinib and gilteritinb showing steric clash with Phe 830. (G) A model of FLT3 active kinase with a stable hydrophobic spine showing the binding of momelotinib and gilteritinb which provides an explanation why both of these inhibitors favor active kinase conformation for binding.

### Momelotinib effectively suppresses growth-factor mediated intrinsic-resistance

Recent studies demonstrated that hematopoietic cytokines GMCSF, IL-3, and FLT3-ligand (FL) promote resistance to FLT3 inhibitors ^20,22^. Expression of FL has been shown to increase during TKI therapy, which conferred resistance to FLT3 inhibitors. Activation of FLT3-WT signaling by FL but not FLT3-ITD was shown to drive the TKI resistance^23,26^. Therefore, we tested the inhibitory activity of momelotinib, quizartinib, and gilteritinib on BaF3 cells expressing FLT3-WT and FLT3-ITD. The addition of FL ligand (50 ng/ml) in BaF3-FLT3-WT cells conferred resistance to quizartinib (>100-fold change in IC50 compared to FLT3-ITD without FL) and gilteritinib (>7-fold increase in IC50) (**Figure 4 A** and **B**). As expected, FLT3-ITD cells did not show any significant change in IC50 values to both quizartinib and gilteritinib in the presence of FL (**Figure 4A** and **B**). In contrast, FLT3-ITD cells treated with FL showed hypersensitivity to momelotinib (2.3-fold reduction in IC50 values compared to non-FL treated cells) while FLT3-WT cells in the presence of FL showed modest increase in IC50 but less than the FLT3-ITD cells without FL (**Figure 4C**). These data suggest that momelotinib, unlike quizartinib and gilteritinib, will suppress the FL mediated resistance.

**Figure 4.**
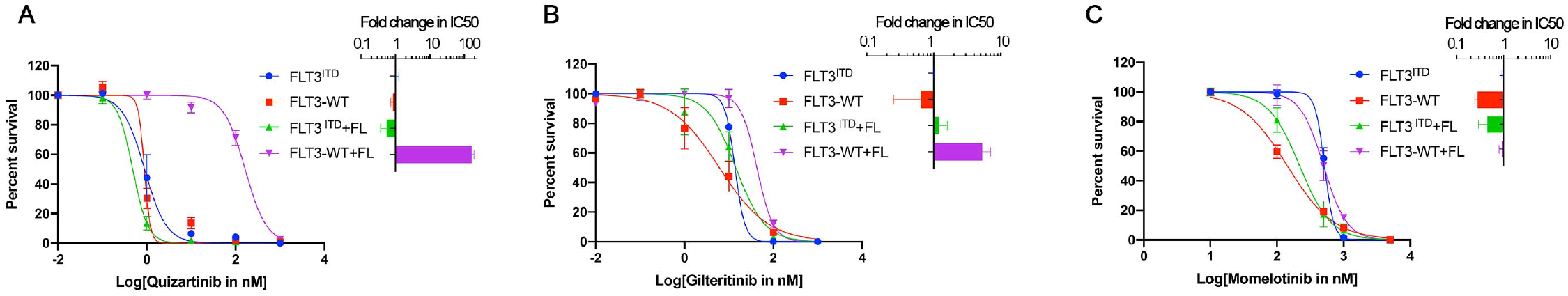
Momelotinib efficiently suppresses the resistance mediated by FLT3 ligand (FL). Sigmoidal curve showing the proliferation of BaF3-FLT3-WT and BaF3-FLT3-ITD in the absence and presence of 50 ng/ml FL treated with increasing concentrations of quizartinib (**A**), gilteritinib (**B**) and momelotinib (**C**). Fold difference in the IC50 values normalized to FLT3-ITD are presented as a bar graph to the right at logarithmic scale. Note, FLT3-WT treated with FL confer significant resistance to quizartinib and gilteritinib while momelotinib fully suppresses the resistance.

A recent study showed that hematopoietic cytokines (GM-CSF and IL3) abrogates FLT3 TKI response by activating JAK2 signaling^22^. Therefore, combinatorial targeting of both FLT3 and JAK2 using a combination of ruxolitinib and gilteritinib showed better efficacy than gilteritinib alone in both in vitro and in a mouse model of leukemia^22^. To test this, we performed a dose-dependent cell proliferation assay to determine the efficacy of momelotinib, quizartinib, and gilteritinib in MV4-11 and Molm13 cells with GM-CSF and IL-3. Both GMC-CSF and IL-3 conferred resistance to quizartinib and gilteritinib (**Supplementary Figure 2**). As expected, momelotinib completely suppressed the GM-CSF and IL-3 mediated resistance (**Supplementary Figure 2**). These data provide evidence that momelotinib is a dual JAK2/FLT3 inhibitor that efficiently suppresses the resistance conferred by FL, GM-CSF, and IL-3.

### Momelotinib treatment reduces leukemic burden in vivo

Preclinical efficacy of FLT3 inhibitors on human leukemia is preferably determined in mouse xenografts using NSG mice. However, a recent study recommends using NSGS mice for accurate *in vivo* modeling as hematopoietic cytokines abrogate TKI response^22^. Therefore, we used NSGS mice to test the efficacy of momelotinib. Mice were transplanted with one million human AML cells Molm13 and MV4-11 expressing luciferase and cherry. Treatment with momelotinib alone (100 mg/kg daily) extended 8-12 days of survival compared to vehicle treated mice (**Figure 5**). Likewise, the leukemic burden from the peripheral blood and BM (deceased or sacked) determined by cherry positive cells showed significant reduction compared to vehicle treated mice. These data validate our finding that momelotinib is an effective FLT3 inhibitor and suppresses the resistance mediated by hematopoietic cytokines.

**Figure 5.**
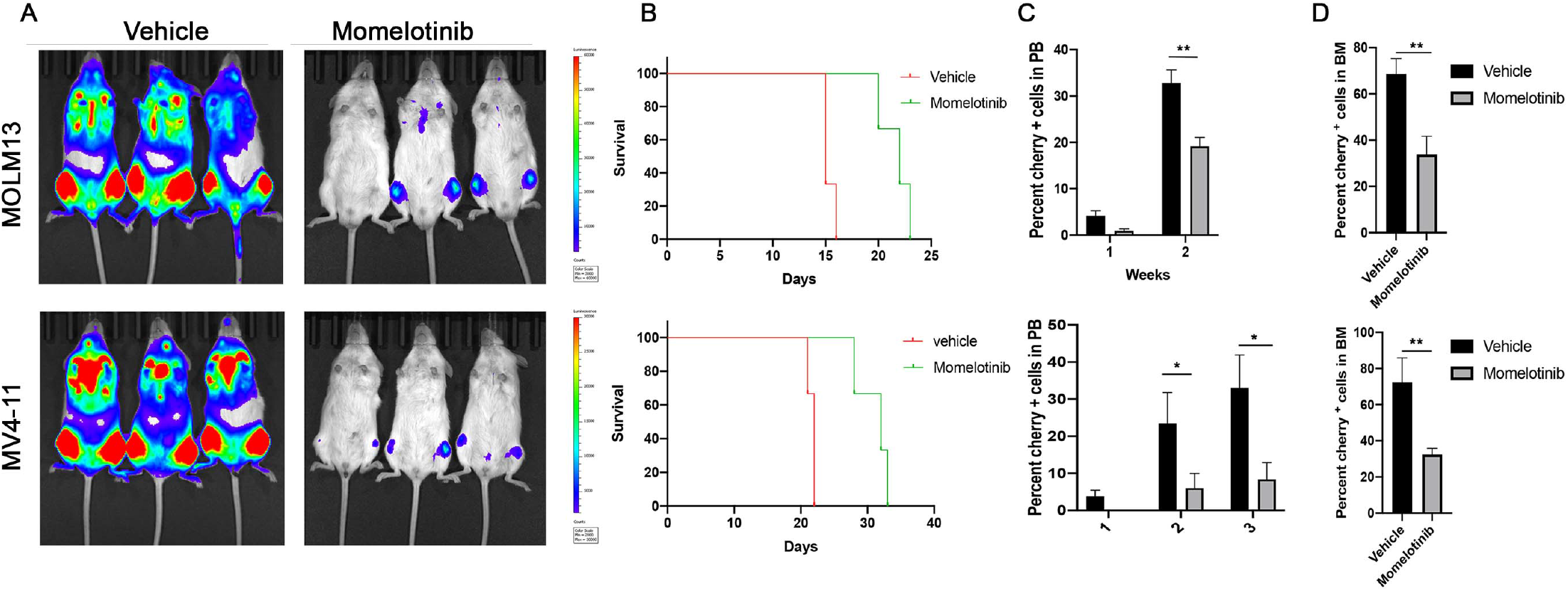
Momelotinib treatment decreases leukemic burden *in-vivo* and prolongs the survival. (**A**) Luminescence imaging of NSGS mice transplanted with MOLM13 (top panel) and MV4-11 (bottom panel) after two weeks of transplantation. One million MOLM13-luciferase-cherry (top panel) and MV4-11-luciferase-cherry (bottom panel) cells were transplanted by tail vein injection in NSGS mice. Momelotinib treatment (10 mg/kg/daily in PBS by i.p injection) was started after two days of transplantation. Control mice were injected with vehicle, PBS. (**B**) shown are survival curve of mice transplanted with MOLM13 (Top panel) and MV4-11 cells (bottom panel). (**C**) Bar graph showing the percentage of leukemic cells determine by cherry positive cells using FACS in mice transplanted with MOLM13 (top panel) and MV4-11 cell (bottom panel). (**D**) Bar graph showing the percentage of leukemic cells in bonemarrow determined by cherry positive cells using FACS in mouse recipients of MOLM13 (top panel) and MV4-11 cells (bottom panel). Error bars represent SEM. \**P* < .05, and \*\**P* < .01.

## Discussion

Several FLT3 inhibitors over the last two decades have been clinically evaluated to treat AML. Three of these, midostaurin, quizartinib, and gilteritinib, have been approved for clinical use in Japan and USA^26^. However, unlike TKI response in CML, the clinical efficacy of FLT3 inhibitors is hindered with both intrinsic and acquired secondary resistance. Intrinsic resistance is predominantly mediated by growth-factor signaling (GM-CSF, IL3, and FLT3 ligand)^20,22^, and differing inhibitory activities of FLT3 inhibitors (especially type-II) against FLT3 mutants (FLT3-ITD and FLT3-TKD)^11,26^. FLT3 type-I inhibitors are equally active on both FLT3-ITD and FLT3-TKD mutants. Secondary resistance is mediated by both on-target (emergence of secondary resistance mutations in FLT3) and off-target mechanism, where leukemic cells activate other signaling pathways for survival and proliferation. Mutations in MAPK pathway genes (NRAS, KRAS, and PTPN11)^17^ and epigenetic modifiers (TET2, IDH1/2, and DNMT3A) ^18^ are commonly observed in off-target mediated resistance. Intrinsic resistance is the root cause of developing acquired resistance as they serve as a reservoir to develop adaptive mechanisms. Momelotinib is a selective JAK2 inhibitor that has been recently approved to treat Myelofibrosis. In this study, we identified that momelotinib is a type-I dual JAK2/FLT3 inhibitor. While the activity of momelotinib was noted against FLT3-ITD positive Molm13 AML cells, its inhibitory activity was ascribed to the off-target effect^24^. Herein we describe the efficacy of momelotinib on FLT3-ITD driven AML in preclinical mouse models and provide a rationale for clinical trials to suppress both intrinsic and acquired resistance to FLT3 inhibitors.

Kinase inhibitors are classified as type-I and type-II inhibitors based on their binding to the ATP-site^37,38^. Inhibitors binding to an inactive conformation are classified as type-II inhibitors, while those preferably binding to active conformation but can also bind to inactive conformations are called type-I inhibitors^37^. The emergence of drug resistant mutations against type-II inhibitors are more frequent than with type-I inhibitors^39,40^. Mutations activating the kinase enzymatic activity or destabilizing the inactive conformation are frequently observed as resistant conferring mutations in relapsed patients treated with type-II inhibitors^12,39,40^. Both quizartinib and sorafenib are type-II inhibitors^41^. Therefore, patients harboring mutations in the activation loop (FLT3-TKD, at residue D835) are intrinsically resistant, and the selection of resistant mutations from the activation loop is commonly observed in patients treated with type-II inhibitors^12,13^. In contrast, type-I inhibitors efficiently suppress the activation loop mutants as their binding is not constrained by activation loop dynamics. The efficacy of momelotinib against the activation loop mutants resistant to type-II inhibitors confirms the *in silico* docking analysis that it is likely a type-I inhibitor. The kinase active site has a number of moving structural modules to be placed precisely for enzymatic activity. The activation loop is one of the most dynamic structures during catalysis and begins with the conserved DFG motif, which adopts two major conformations representing active and inactive state^37^. DFG-in conformation represents the active state where aspartate residue moved in facing the active site and coordinates with β-phosphate of ATP through a magnesium ion, and Phe residue moves away from the ATP site to stabilize the hydrophobic spine by coordinating with C-helix and catalytic motif HRD (Figure 3G). DFG-out conformation represents the inactive state where Asp and Phe swap their positions resulting to a dismantled hydrophobic spine and broken interaction between aspartate and ATP. Mutations in the activation loop increase its flexibility, which increases the equilibrium favoring DFG-in conformation (active state)^42,43^. Depending on types of substitution, some mutations will be more activating than the others. For instance, different amino acid substitutions at Asp 835 displayed altered inhibitor sensitivity that correlates with enzymatic activation, such that most active mutant conferred greater resistance^44^. Our docking prediction revealed that momelotinib binds to the active conformation, which explains its activity against the activation loop mutants. However, momelotinib failed to inhibit the gatekeeper mutant F691L though it does not show any physical interaction. The gatekeeper mutation F691L confers resistance to all clinically used FLT3 inhibitors by seemingly causing a steric blockade to inhibitor binding, which, nonetheless, is true for resistance to quizartinib and midostaurin. However, we did not observe any steric hindrance posed by F691L mutation to either momelotinib or gilteritinib suggesting alternate mechanism for drug resistance. Given that the gatekeeper residue governs conformational state by modulating the assembly and destabilization of the hydrophobic spine, we hypothesize that F691L mutant stabilizes an intermediate kinase conformation to which type-I inhibitors have lower affinity. Earlier, we have shown that substitutions of gatekeeper residue with a bulkier hydrophobic residue (such as isoleucine or methionine) in oncogenic kinases (ABL, PDGFR, SRC and EGFR) activates the kinase by stabilizing the hydrophobic spine while substitution with smaller hydrophobic residue (alanine) destabilizes the spine^42^. Likewise, substitution of smaller hydrophobic residues at the gatekeeper residue in ERK2 (Ala for Glu 130) create a cavity within the hydrophobic cluster that destabilizes the autoinhibited state and increases the flexibility of the DFG motif and activation loop^43^. Increased flexibility of the activation segment is expected to favor activation loop phosphorylation, which stabilizes the active conformation. These observations suggest that substitution of a less bulky residue, Leu, for Phe 691 may destabilize the hydrophobic spine that favors an inactive conformation while increased flexibility of DFG motif and activation loop will shift the equilibrium towards the active state conformation, which may likely result in stabilizing an intermediate conformation. Our *in vitro* cell proliferation data from the compound FLT3-ITD mutation (FLT3-ITD-F691L/Y842H) supports this notion. For instance, we observed a significant reduction in momelotinib resistance against the compound mutant, F691L+Y842H, compared to F691L alone while FLT3ITD-F691L/D835Y did not show any significant change in IC50. This increased inhibitor sensitivity of FLT3-ITD-F691L/Y842H correlates with elevated auto-phosphorylation compared to resistant compound mutant (FLT3-ITD-F691L/D835Y), which suggests that stabilization of active state conformation would restore momelotinib sensitivity against the gatekeeper mutant. Altogether these data support that momelotinib binds to an active and open conformation of the kinase.

FLT3 TKIs show differential responses to leukemic cells. For instance, FLT3 inhibitors induce apoptosis in circulating blasts but promote differentiation of leukemic blasts from the bone marrow. This suggests that BM niche is a potential source of intrinsic resistance resulting in leukemic persistence and minimal residual disease. These surviving leukemic cells serve as a reservoir to develop resistance. Hematopoietic cytokines and growth factors (FL and FGF2)^20,21^ released from the BM stroma/niche abrogate oncogene-dependence resulting in TKI resistance. Notably, FL, whose expression is reported to increase during chemotherapy abrogates TKI response by activating canonical FLT3 wild-type signaling^20,23^. However, it is not clear how activation of FLT3-WT confers resistance. Nonetheless, an altered signaling network engaged by FLT3-WT supporting survival during TKI therapy is postulated possibly exploiting the cell-intrinsic inflammatory-related pathways for survival ^45,46^. Likewise, cytokine-induced activation of Jak2 signaling confers resistance^22^. In addition, high levels of phospho-JAK2 are associated with adverse clinical outcomes in AML, which seemingly stems from elevated inflammatory cytokines. Therefore, concomitant targeting of FLT3 and cytokine signaling using, for instance, JAK2 inhibitor (ruxolitinib) showed an improved response in preclinical mouse models of AML. We show that momelotinib, by targeting both FLT3 and JAK2, suppresses both cytokines (GMCSF and IL3) and FL mediated intrinsic resistance. We anticipate that momelotinib will be more effective in suppressing the intrinsic resistance than the currently used FLT3 clinical inhibitors.

The majority of FLT3 inhibitors exhibit cross-reactivity with c-KIT. Concomitant inhibition of FLT3 and c-KIT (even partial inhibition) shows poor hematopoietic recovery during treatment, resulting in myelosuppression. For instance, quizartinib treatment showed a >50% response rate in FLT3-ITD positive AML cases. However, these responses were associated with incomplete blood recovery (CRi) due to inhibition of c-KIT. In contrast, gilteritinb, which does not inhibit c-KIT showed superior complete remission (CR) with improved blood recovery. Our data demonstrate that momelotinib, like gilteritinb, does not inhibit c-KIT suggesting that it will not be myelosuppressive (**Supplementary Fig 3**). Recent clinical studies showed that momelotinib treatment is not myelosuppressive rather it alleviates anemic symptoms in most MF patients and did not show any significant cardiotoxicity^25,27^. Perhaps more interestingly, unlike gilteritinib, momelotinib inhibits the expression of MYC (**Supplementary Fig 4**). Elevated MYC expression is one of the hallmarks of cancer, including AML. So far, targeting MYC has been a clinical challenge. Nonetheless, suppression of MYC expression by BRD4 inhibitors in AML are showing promising results in preclinical AML models^29,30^. Given that the momelotinib inhibits MYC expression, we anticipate that it will be more effective in suppressing the disease than the currently used type-1 FLT3 inhibitors.

Altogether, momelotinib has several advantages over currently used FLT3 TKIs. Poor efficacy of type-II FLT3 inhibitors is associated with; frequent emergence of drug-resistant mutants, myelosuppression due to off-target c-KIT inhibition, and loss of therapeutic response under cytokine signaling. We demonstrate that momelotinib is a type-1 dual JAK2/FLT3 Inhibitor, effectively suppresses the resistance mediated by activation loop mutants and growth factor signaling. Besides, we do not anticipate that it will be myelosuppressive as it lacks activity against c-KIT. Severe cases of myelosuppression are not reported from the clinical studies performed with momelotinib supports our finding. Moreover, its activity in suppressing MYC expression suggests that it may provide a better therapeutic outcome. Altogether these data support that momelotinib will induce a deep and durable clinical response in AML by suppressing intrinsic and acquired resistance. Thus, warrants its clinical evaluation in AML patients.

## Acknowledgements

This study was supported by grants to MA from the National Cancer Institutes at NIH (RO1CA211594) and (RO1CA250516). MA is a recipient of the bridge award from the American society of hematology (ASH). Authors are thankful to Erika Huber for performing initial cell proliferation assays and western blotting.

## Author Contribution

M.A., M.K. and Z.K performed all experimentation and analyses. M.A., M.K., Z.K, and M.A designed all experiments. DTS provided critical reagents and guided in data analyses. M.A. M.K, and M.A wrote the manuscript.

