## Supplementary Figure legend for "Momelotinib is a highly potent inhibitor of FLT3-mutant AML"

**Supplementary Figure legends:**

**Supplementary Figure 1.  Ruxolitinib does not inhibit FLT3 kinase**

(**A**) Sigmoidal plot showing the survival of BaF3, BaF3-FLT3-WT, and BaF3-FLT3-ITD treated with increasing concentrations of ruxolitinib for 72 hrs. (**B**) Immunoblot showing the phospho-FLT3, and phosphor-STAT5 in cells treated with increasing concentrations of ruxolitinib for six hours. Unlike momelotinib, ruxolitinib does not inhibit FLT3-WT or FLT3-ITD. (**C** and **D**) Sigmoidal curve showing the viability of AML cells, MOLM13, MV4-11 and K562 treated with quizartinib (C) and gilteritinib (D).

**Supplementary Figure 2**. **Momelotinib efficiently suppresses the resistance mediated by hematopoietic cytokines GM-CSF and IL3.**

(**A-C**) Sigmoidal curve showing the viability of MOLM13 cells grown with GM-CSF (50 ng/ml) or IL3 (20ng/ml) and treated with increasing concentrations of quizartinib (A), gilteritinb (B) and momelotinib (C). (D-E) Sigmoidal curve showing the survival of MV4-11 cells in the presence of GM-CSF (50 ng/ml) or IL3 (20ng/ml) treated with increasing concentrations of quizartinib (D), gilteritinb (E) and momelotinib (F). Fold difference in the IC_50_ values normalized to FLT3-ITD are shown on the right side as a bar graph at logarithmic scale.

**Supplementary Figure 3**. **Momelotinib does not inhibit c-KIT receptor.**

Sigmoidal curve showing the viability of BaF3, BaF3-KIT-D816V (TKI sensitive mutant) and BaF3-D816V/T681I (TKI resistant variant due to mutation of gatekeeper residue, T681I) treated with dasatinib (**A**), quizartinib (**B**), gilteritinb (**C**) and momelotinib (**D**). Note, well know KIT inhibitors dasatinib and quizartinib efficiently suppress the KIT-D816V mutant but failed to inhibit the gatekeeper variant T681I suggesting that the observed inhibition is due to on-target inhibition. In contrast, momelotinib like gilteritinib failed to inhibit the KIT expressing cells suggesting that it does not have activity against the KIT receptor.

**Supplementary Figure 4.  Momelotinib suppresses the expression of MYC**

Immunoblot analysis showing the expression of MYC in MOLM13 (left panel) and MV4-11 cells (right panel) treated with momelotinib and gilteritinib. Concentration of inhibitors are indicated above the lanes. Total cell extracts were probed with anti-Myc antibody and b-actin antibody. Note, unlike gilteritinib, momelotinib efficiently suppress the expression of Myc suggesting that it might induce deeper clinical response
