## Supplementary figures and images for "Momelotinib is a highly potent inhibitor of FLT3-mutant AML"

### Supplementary Figure 1

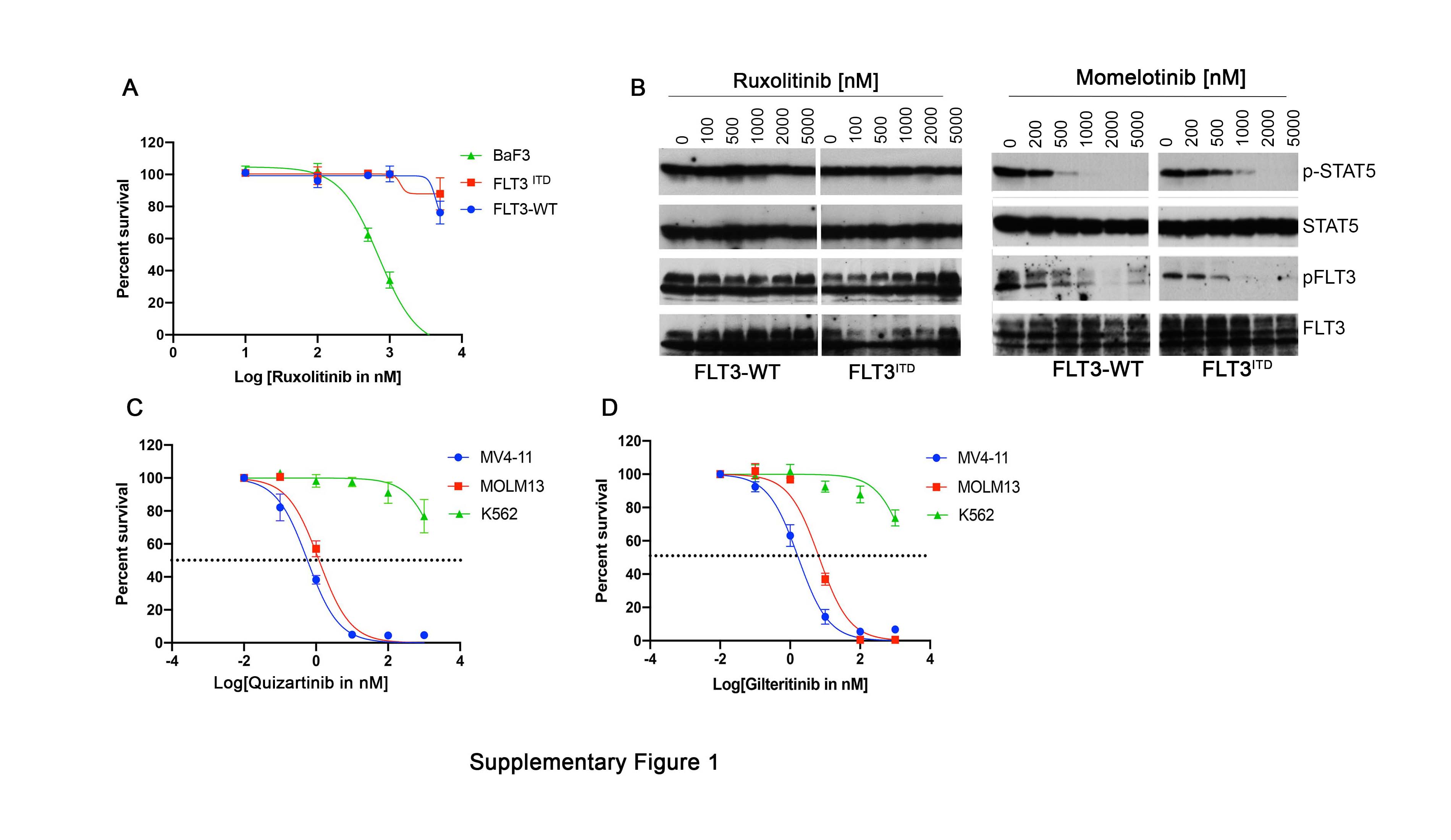

### Supplementary Figure 2

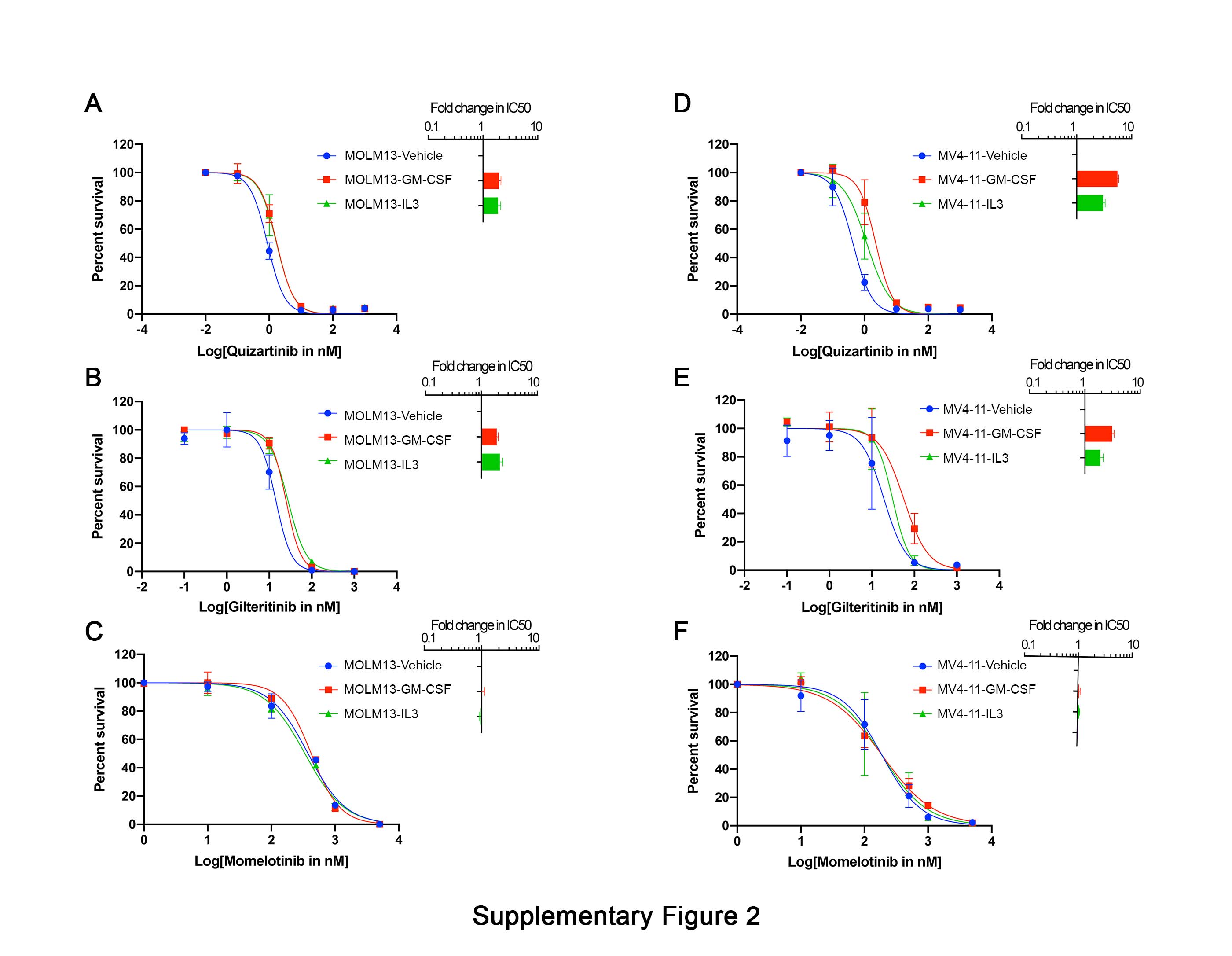

### Supplementary Figure 3

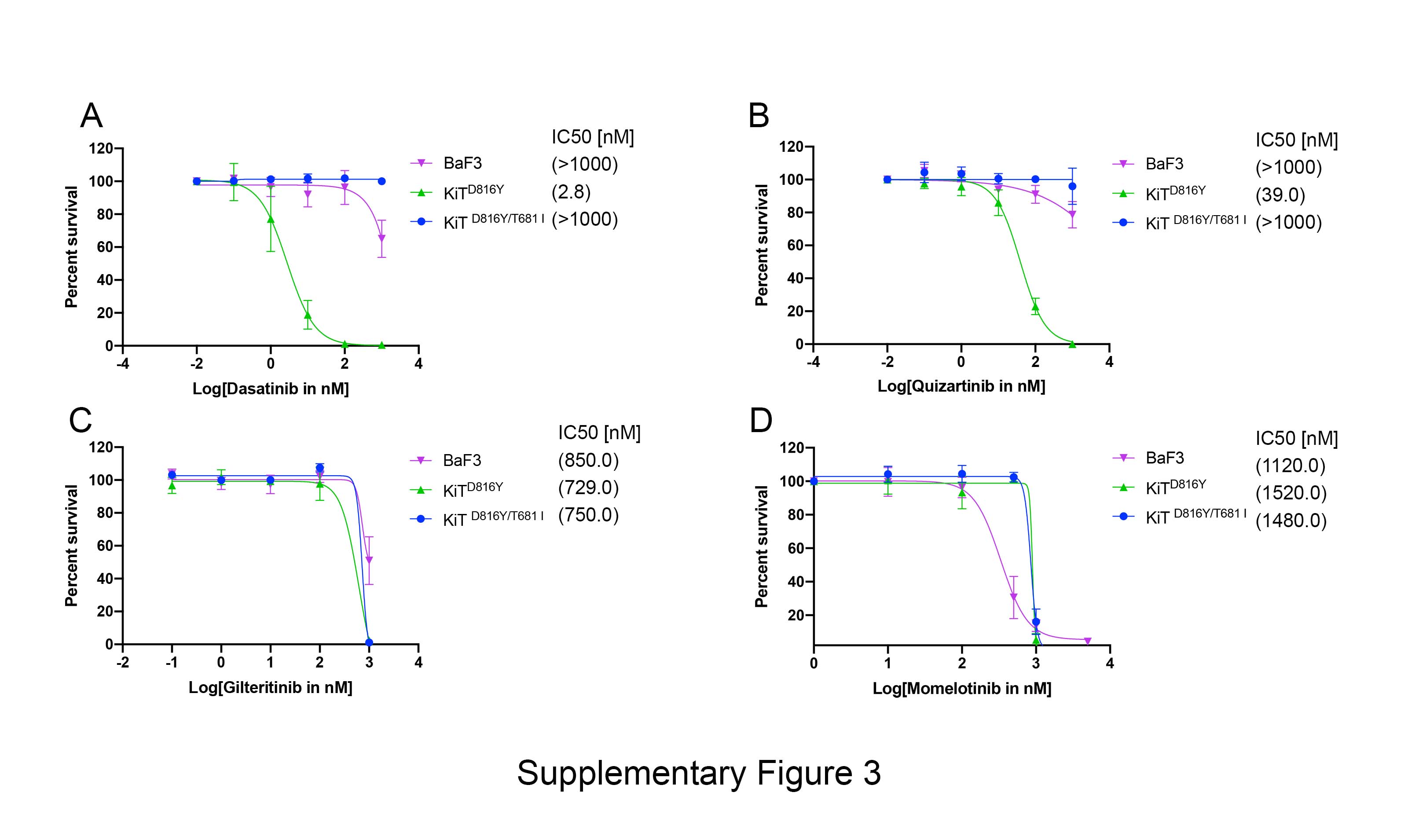

### Supplementary Figure 4

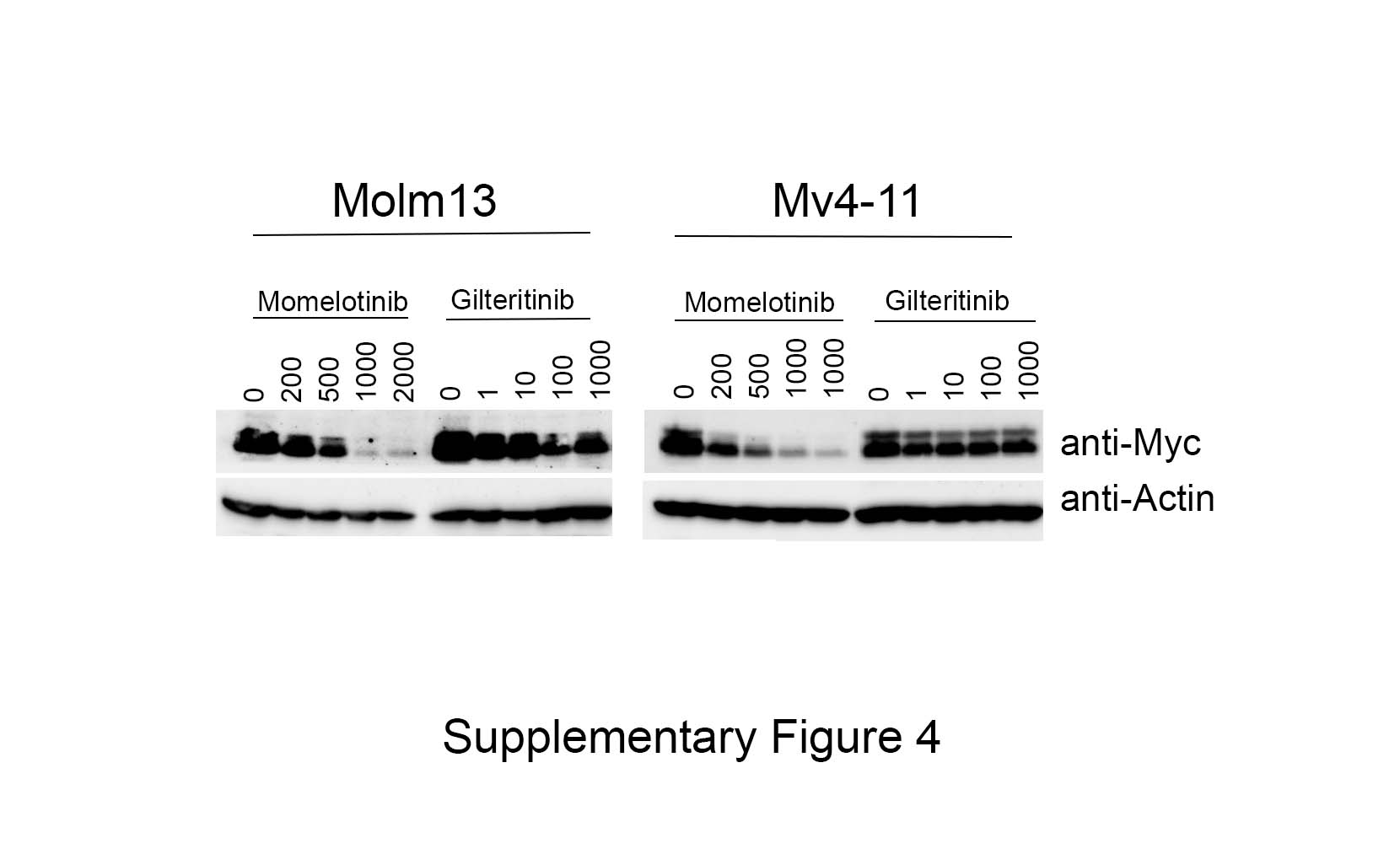
